# Sparking “The BBC Four Pandemic”: Leveraging citizen science and mobile phones to model the spread of disease

**DOI:** 10.1101/479154

**Authors:** Stephen M. Kissler, Petra Klepac, Maria Tang, Andrew J.K. Conlan, Julia R. Gog

## Abstract

The nexus of mobile technology, mass media, and public engagement is opening new opportunities for research into the human behaviours relevant to the spread of disease. On 22 March 2018, the British Broadcasting Corporation (BBC) released the documentary “Contagion! The BBC Four Pandemic” to describe the science behind pandemic preparedness in the UK. The authors of this article were responsible for producing a mathematical simulation for that documentary of how a highly contagious respiratory pathogen might spread across the UK. According to the documentary narrative, the ‘outbreak’ begins in the town of Haslemere, England. To ground the simulation in true human interaction patterns, a three-day citizen science experiment was conducted during which the pairwise distances between 469 volunteers in Haslemere were tracked continuously using a mobile phone app. Here, we offer a scientific companion to the documentary in which we describe the methods behind our simulation and release the pairwise interpersonal distance dataset. We discuss salient features of the dataset, including daily patterns in the clustering and volatility of interpersonal interactions. Our epidemiological analysis of the simulated Haslemere outbreak serves as a springboard to discuss scientific opportunities opened by the Haslemere dataset and others like it. We believe that the Haslemere dataset will productively challenge current strategies for incorporating population structure into disease transmission models, and hope that it will inspire the collection and analysis of other similar datasets in the future.

## 1 Introduction

On 22 March 2018, the television channel BBC Four released “Contagion! The BBC Four Pandemic” [1], a documentary on influenza epidemiology to mark the centenary of the 1918 Spanish Influenza pandemic. The documentary explores how a highly contagious new strain of influenza could spread across the United Kingdom using a simulated outbreak based on mathematical models developed by the authors of this article. According to the documentary narrative, a simulated outbreak originates in Haslemere, a town in Surrey, in the south of England, and then continues on to infect the rest of the UK. In this scientific companion to the documentary, we present a novel dataset that captures the interpersonal distances between 469 volunteers from Haslemere over three days, and we describe a mathematical model that incorporates these data, used to simulate the “Haslemere outbreak” featured in the television programme. A description of the UK-wide model is given by Klepac *et al*. (2018) [2].

The production of the documentary provided an ideal opportunity to engage the public in collecting scientifically valuable data on interpersonal contact patterns. Close-proximity interactions are the main avenue by which many infections spread, and yet we lack a detailed understanding of when, with whom, and with what frequencies and durations these interactions normally occur. This is due in part to a lack of adequate data, which can in turn be traced to technological limitations and to issues with privacy. Wearable proximity sensors can provide detailed measurements of interpersonal distances [3, 4], but these devices can be expensive and require substantial oversight in their deployment and collection, limiting the scope of studies that rely on them. Mobile phones are an attractive alternative, since many people carry them and they feature a range of sensing technologies that can both identify nearby devices and measure GPS location. Bluetooth, for example, can identify nearby devices with a simple ‘is/is not within signal range’ dichotomy [5, 6], but is unreliable for measuring actual distances. Mobile phone GPS capabilities, on the other hand, can pinpoint a user’s location using various strategies including satellite-mediated GPS, cellular tower triangulation, and wifi connection. These GPS tags can then be used to measure the distances between phones. GPS accuracy is imperfect [7], however, and the interpersonal distances measured by GPS cannot take physical barriers (e.g. walls) between the users into account. A range of studies have used mobile phones to study human mobility patterns [3, 4, 5, 6, 7, 8, 9, 10, 11, 12, 13, 14], but due in part to privacy concerns the majority of these only consider bulk movements, obscuring the fine-scale interaction patterns that underpin the spread of disease. The study by Palmer *et al*. (2013) [7] most closely resembles the one presented here, though the scope is global rather than within a single town, covers fewer participants, and the dataset is not publicly available.

Our understanding of the nature of interpersonal interactions is also limited by a lack of adequate mathematical vocabulary to describe their intricate, often ‘fleeting’ structure [7, 15]. Networks are currently a popular way of incorporating population structure into models of infectious disease transmission [16, 17, 18, 19, 20, 21, 22, 23]. Static, unweighted networks are particularly amenable to mathematical analysis, and have featured in a range of disease transmission models [22, 24, 23, 25, 26, 27, 28]. However, there is a growing body of evidence that suggests that static networks may not adequately capture information about the frequency, proximity, and timing of interpersonal encounters, critical to the spread of disease. Interest in dynamic networks is increasing, both from theoretical [15, 29, 30] and empirical [31, 32, 33, 34] standpoints. Datasets like the one presented here may help ground future dynamic network models in reality, and may help reveal what information is lost when the the rich dynamics of human interactions are approximated by a static network.

The Haslemere dataset enhances our capability to examine disease transmission and human behaviour in general. While we focus primarily on the work carried out for the BBC Four documentary, we also briefly discuss other aspects of disease transmission that might be examined using the Haslemere dataset and others like it. Following Lloyd-Smith *et al*. (2005) [35], we characterise the superspreaders of the Haslemere “outbreak”, and we examine the challenges associated with empirically estimating the basic reproduction number *R*_0_ for outbreaks in structured populations. We conclude with a discussion of lessons learned and open questions that might be addressed using datasets like the one presented here.

## 2 Description of the Haslemere dataset

The Haslemere dataset (Supplemental Data 1) consists of the pairwise distances of up to 1m resolution between 469 volunteers from Haslemere, England, at five-minute intervals over three consecutive days (Thursday 12 Oct – Saturday 14 Oct, 2017), excluding the hours between 11pm and 7am. The volunteers constitute a convenience sample of 4.2% of the total population of Haslemere, according to the 2011 UK census [36]. Participants downloaded the *BBC Pandemic* mobile phone app and then went about their daily business, with the app running in the background. Ethical guidelines required that the study be restricted to volunteers at least 16 years of age, or at least 13 years of age with parental consent. The pairwise distances between volunteers were calculated using Haversine formula for great-circle geographic distance and are based on the most accurate GPS coordinates that the volunteers’ mobile phones could provide. Full details on the study site, recruitment, data collection, and post-processing are provided in the SI.

Fig. 1 depicts the pairwise encounters within 20m for a subset of the volunteers as a network, where each edge is coloured according to the quarter-day in which the encounter occurred [37]. This illustrates the heterogeneity and temporal complexity of the data, with some clustered individuals who encounter each other frequently, and other individuals who provide infrequent links between distant parts of the network. Relatedly, Movie S1 depicts the pairwise interpersonal probabilities of infection for a subset of volunteers under the Haslemere outbreak transmission model (see SI), at four different temporal resolutions. The video illustrates how some connections persist over time, while other dynamic elements are lost when the network is temporally integrated.

**Figure 1:**
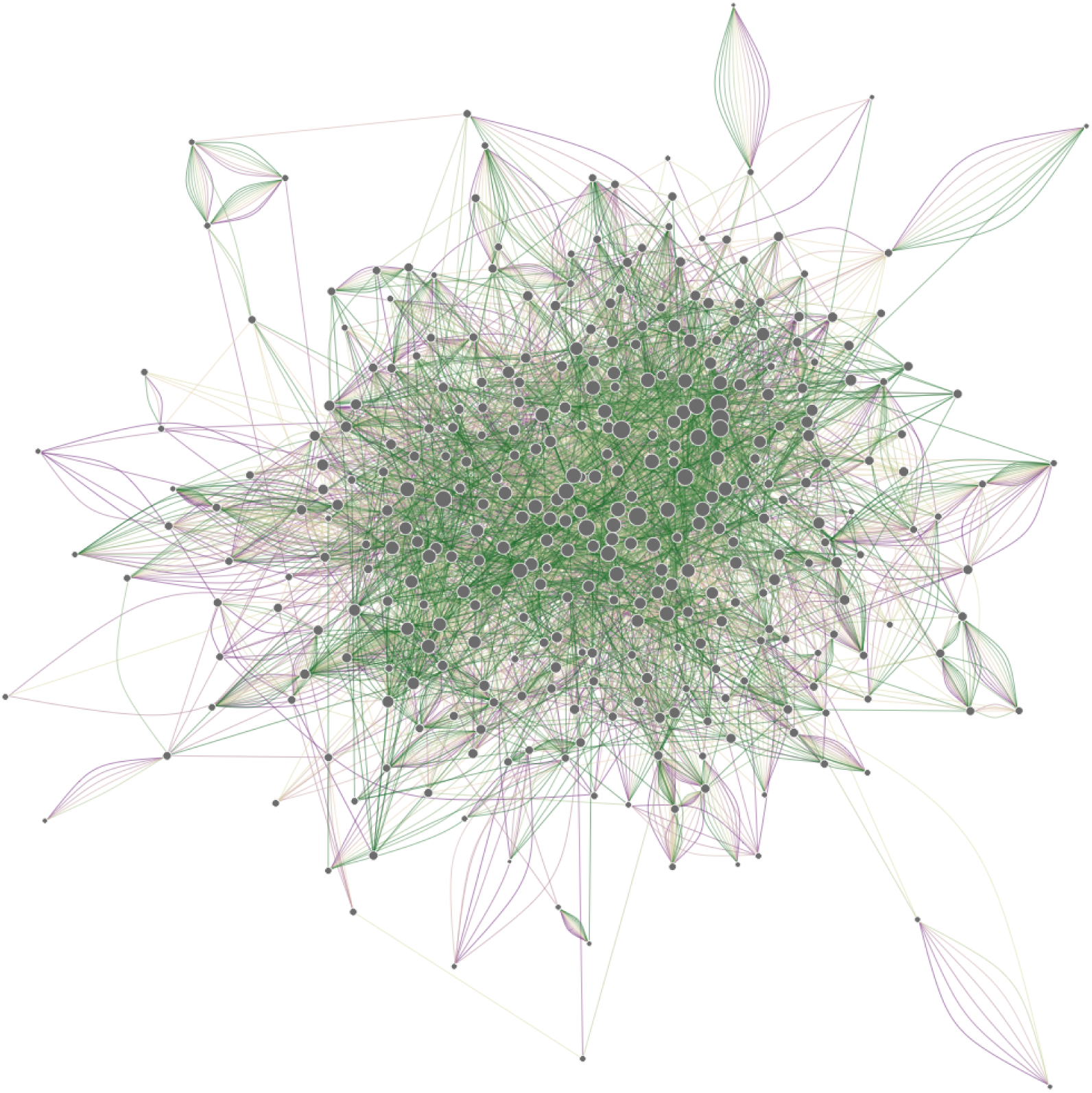
Network depicting pairwise encounters over time for the 75% of individuals in the Haslemere dataset who spend the greatest amount of time within 20m of another person (i.e. with the most total ‘person-hours’ of contact). An encounter is defined as the first time two individuals come within 20m of one another during some span of time. Each node represents an individual, and each line represents an encounter that occurred during a particular quarter of one day. Quarter-days consist of the hours between 7am–11am, 11am–3pm, 3pm–7pm, and 7pm–11pm. There are twelve quarter-days across the three days of the study, and thus at most twelve lines that can connect any two nodes. Lines are coloured according to the quarter-day in which the encounter occurred, ranging from Thursday Quarter 1 (purple) to Saturday Quarter 4 (green). Node area is proportional to the total number of unique encounters for that individual across all three days (which would be that individual’s degree, if the network were represented without temporal information).

Although the Haslemere dataset is among the most comprehensive of its kind, it is still subject to a number of limitations. Since a substantial fraction of Haslemere residents did not participate in the study, the dataset (and any network representation, such as Fig. 1) is unrealistically sparse. The interaction patterns are also biased by participant age, since children under 13 were not allowed to participate and the rate of smartphone ownership declines substantially in adults over 65 [38]. A literal interpretation of the contact network is therefore inadvisable; instead, the dataset should be understood to represent a proportion of adult encounters, from which general patterns may be identified with some caution. Epidemic models based on the Haslemere dataset should take these limitations into account.

### Interaction patterns by time of day

The Haslemere dataset provides an opportunity to examine how interpersonal interaction patterns within a town vary by time of day. One might expect individuals to have many fleeting interactions during the day, and fewer, more persistent interactions at night. The Haslemere dataset supports this hypothesis. To illustrate this, we calculate the mean degree, mean local clustering coefficient [39], and mean link volatility [40] (Fig. 2) during five-minute network ‘snapshots’ generated from the pairwise Haslemere distance data. For each time point in the dataset, corresponding to a five-minute interval in real time, a network is generated in which each node represents an individual, and a link is defined between two nodes if the corresponding individuals are within some cutoff distance of one another during that five-minute interval. The mean link volatility 1 *– γ*, introduced by Clauset and Eagle (2012) [40], extends the Pearson correlation coefficient to measure the persistence (*γ*) of network links between subsequent snapshots (see Materials and Methods). For all cutoff distances, the mean degree and mean clustering coefficient are highest during the nighttime hours, while the mean link volatility is highest during the day (Fig. 2). This suggests that people tend to form temporally persistent small groups in the evenings, perhaps as families occupying the same household, and tend to have more fleeting interactions during the day with a wider range of people. This contrasts with Clauset and Eagle [40], who find in a study of the interpersonal interactions of students and professors at the Massachusetts Institute of Technology that the mean degree, mean local clustering coefficient, and mean volatility are all highest during the day. This difference may be attributed to the fact that the Haslemere study captures the movements of people across a whole town, and thus represents a more general cross-section of the population.

**Figure 2:**
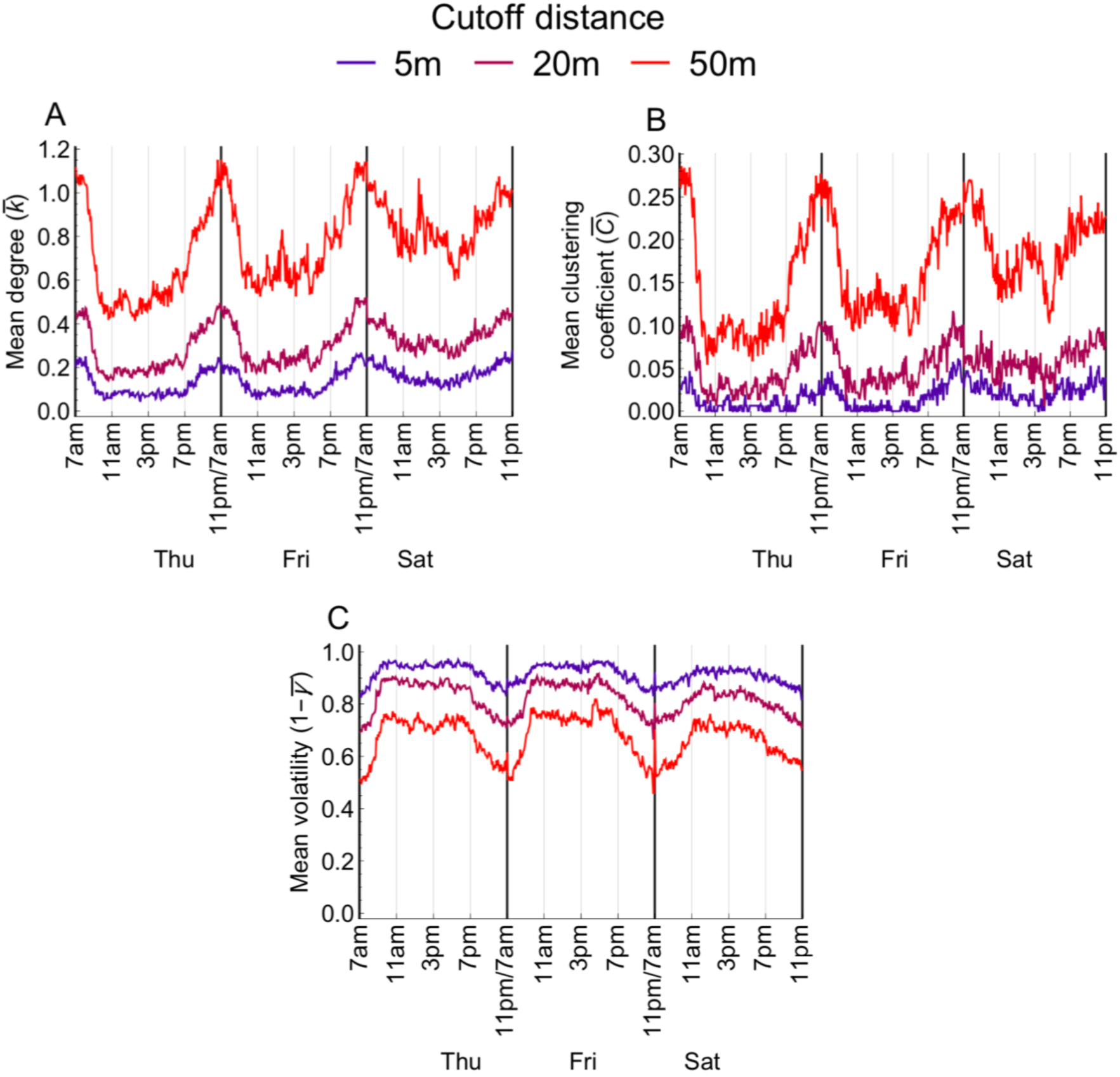
(A) Mean degree, (B) mean local clustering coefficient, and (C) mean link volatility over time for five-minute network ‘snapshots’ generated from the Haslemere dataset. Mean degree and mean local clustering coefficient are highest at night, while the mean link volatility is highest during the day.

These findings suggest that interpersonal interaction patterns might productively be separated into a ‘daytime’ and ‘nighttime’ regime, or possibly into ‘school/work’ and ‘home’. For epidemiological applications, one might be most interested in daytime interactions when describing the spread of a disease transmitted through casual contact (e.g. measles), while one might wish to know more about nighttime contacts when describing the spread of a disease transmitted mainly through persistent contact (e.g. tuberculosis). For still other diseases, some combination of the two will be relevant. We recommend including time of day and/or geographic venue as an element in future interpersonal contact surveys that are intended to support epidemiological research.

## 3 The BBC Pandemic simulation

To generate the simulated outbreak for the BBC documentary, we constructed a minimal susceptible-exposed-infectious (SEI) model that incorporates the Haslemere dataset. Here, we discuss that model and present a brief epidemiological analysis. The Haslemere outbreak simulation was produced in real time during a three-day data collection period. The participants’ GPS locations were sent nightly to the authors of this article to develop and test a disease transmission model. Immediately following the final data pull, we simulated a single outbreak from the model to feature in the documentary. To fit into the programme narrative, the “outbreak” needed to be seeded by the presenter (user 469), and last for a total of three days. Because of this, some scientific liberties were taken: model parameter values were chosen to ensure that a substantial outbreak would occur within the three-day simulation window, requiring an unrealistically high basic reproduction number (*R*_0_) and a short incubation period. Motivation for the chosen model parameter values may be found in the SI. Despite these liberties, this minimal model satisfies the central aim of communicating key aspects of respiratory infectious disease transmission, and can be easily adjusted to account for other scenarios. Full details on model specification are given in the Materials and Methods, and computer code for producing related simulations is given in Supplemental Code 1. All of the following references to the “Haslemere outbreak” refer to our computational simulations; no real illness was tracked in this study.

We used the transmission model to generate a single epidemic simulation, which we call the “the Haslemere outbreak”, to feature in the BBC Four documentary. A total of 405 participants were ‘infected’ in the simulated Haslemere outbreak over the three days (Fig. 3A). Generations of infection were identified by assigning to each infection an immediate ‘parent’, taken to be the geographically nearest infectious individual at the time of infection. While this stretches the underlying theory somewhat – infections are actually caused by the accumulation of forces from nearby infectious individuals, not by a single progenitor – this approach serves to illustrate key transmission routes for this particular simulation. Fig. 3B depicts these generations of infection as a transmission tree. The longest transmission chain consists of 14 infections.

**Figure 3:**
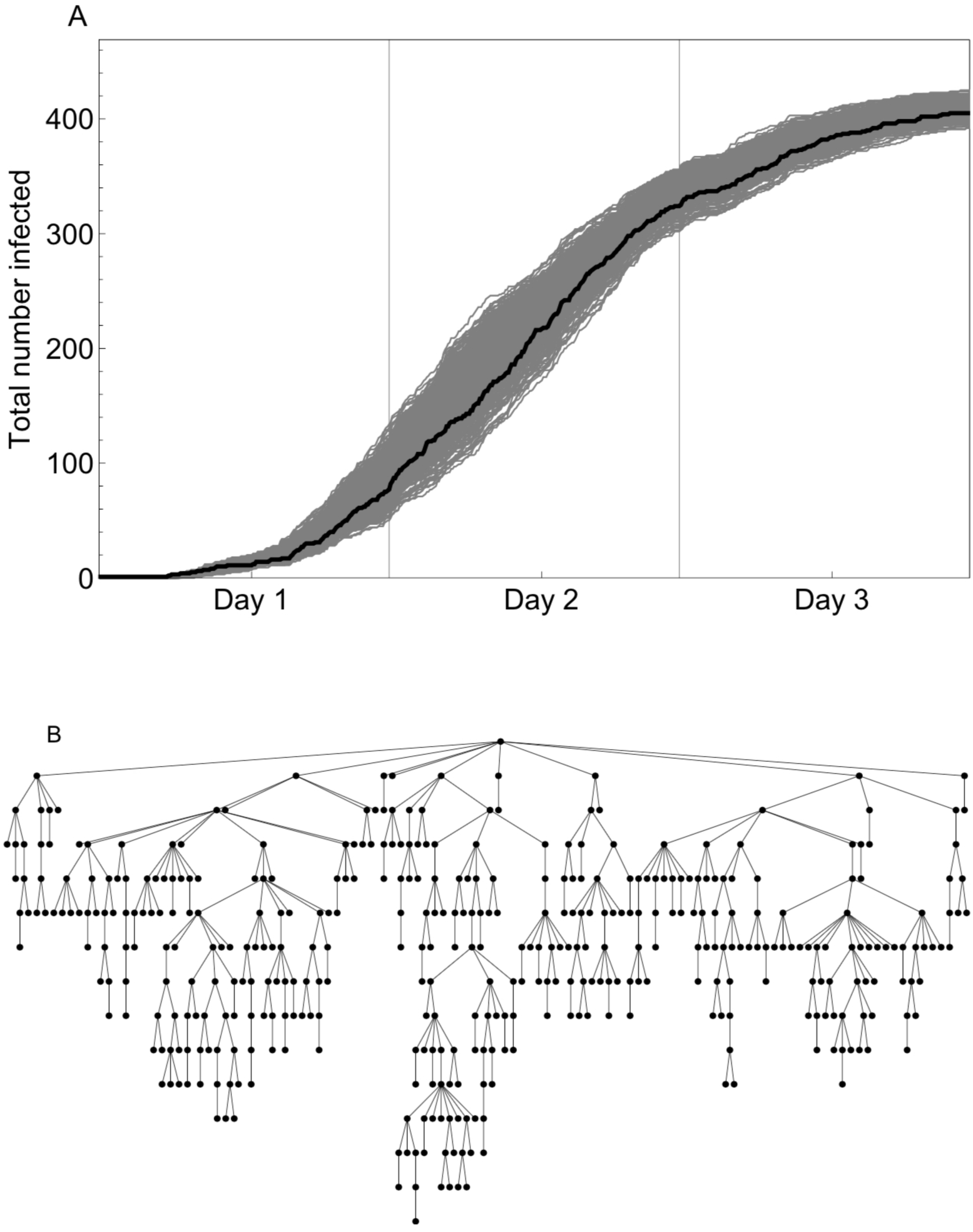
(A) Cumulative number of people infected over time in the Haslemere epidemic (black) and in 500 re-simulations (grey). (B) Tree depicting direct infections in the Haslemere epidemic. Each row represents a new generation of the outbreak, from top to bottom. Note that infections that align on a row did not necessarily occur at the same time, but instead represent “generations” of the outbreak. The point at the root of the tree (top) represents the index case, who caused nine direct secondary infections.

### Identifying superspreaders

Individuals that are responsible for disproportionately many infections during an outbreak are commonly termed “superspreaders”. Superspreaders are natural targets for interventions aimed at reducing disease transmission, but are notoriously difficult to identify [35, 41, 42]. Here, we ask whether the superspreaders of the Haslemere outbreak can be identified using time of infection (are the superspreaders simply those who happened to get infected early in an outbreak?), number of encounters (are the superspreaders those who interact with the greatest number of people?), or a model-based individual reproduction number (are the superspreaders those with the greatest infectious potential?). To identify the superspreaders of the Haslemere outbreak, we use our earlier strategy of tracing each infection to a “most likely” infectious progenitor. By this metric, most (272) individuals caused no infections, but a few caused disproportionately many (Fig. 4A). Lloyd-Smith et al. [35] found that negative binomial distributions tend to fit a wide variety of offspring distributions observed during real outbreaks. If *Z ∼ NegBin*(*r, p*) is the offspring distribution, an *n*th-percentile superspreader is defined as anyone who infects at least *Z*^(*n*)^ others, where *Z*^(*n*)^ is the *n*th percentile of the offspring distribution [35]. For the Haslemere epidemic, the 90th-percentile superspreaders are those who infected at least *Z*^(90)^ = 3 individuals, of whom there were 47.

**Figure 4:**
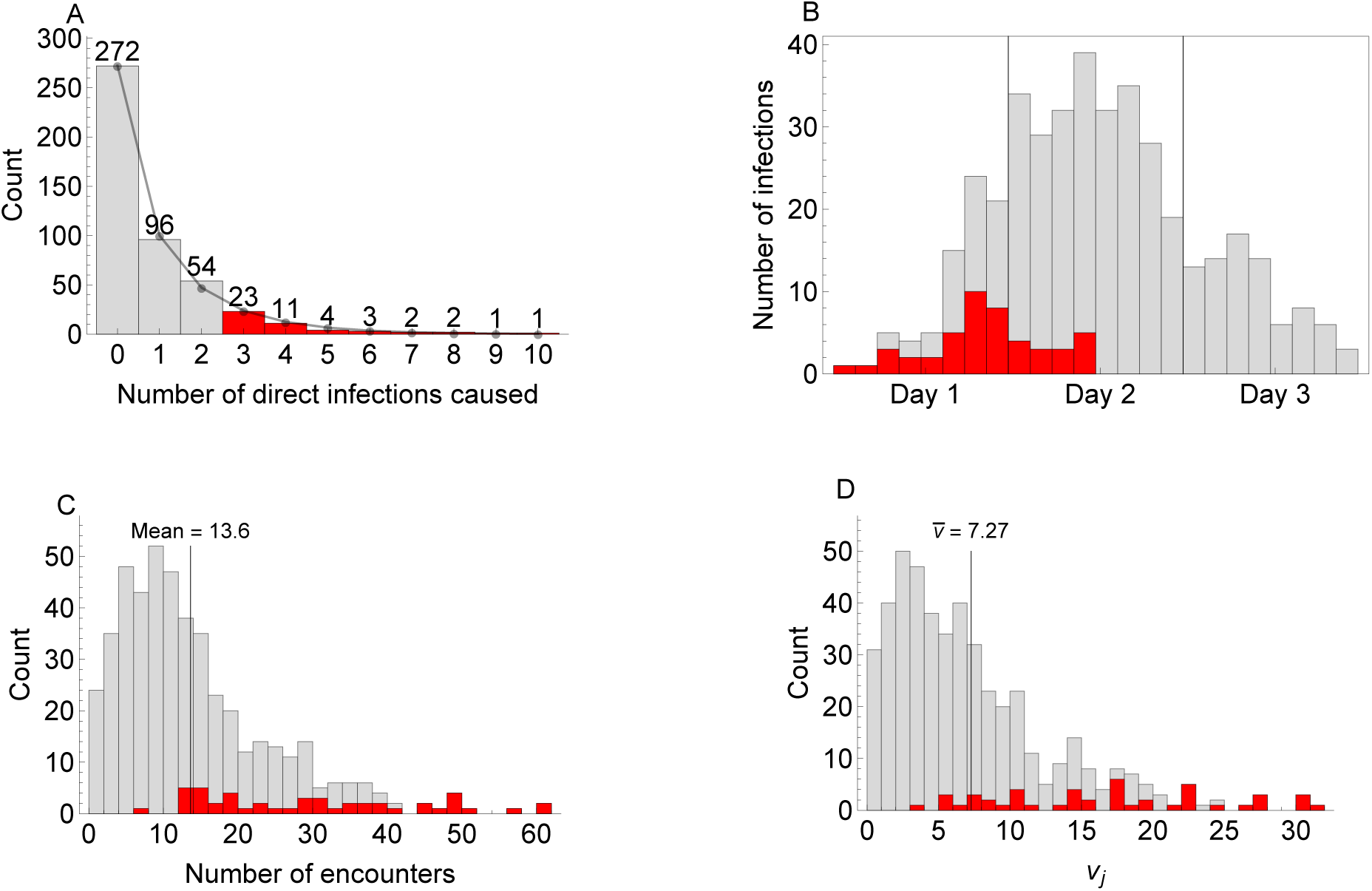
(A) Histogram of the number of direct secondary infections caused by each person in the Haslemere epidemic. The connected points depict the best-fit negative binomial distribution (*r* = 0.640, *p* = 0.426) to the histogram values. The outbreak’s 90th-percentile superspreaders are the 10% of people (*n* = 47) who caused the greatest number of direct secondary infections (3 or more). These are highlighted in red. (B) Histogram of the total number of people infected over time (grey), and the number of the 90th-percentile superspreaders infected over time (red). All superspreaders are infected by the middle of Day 2 of the outbreak. (C) Histogram of the number of unique encounters within 20m for all users (grey) and for the 90th-percentile superspreaders (red). The mean number of unique encounters is 13.6. (D) Histogram of the individual reproduction number *v*_*j*_ for all users (grey) and for the 47 90th-percentile superspreaders (red). The mean individual reproduction number is 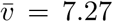, which may also be interpreted as the overall basic reproduction number, *R*_0_.

The superspreaders of the Haslemere outbreak were generally infected early in the epidemic (Fig. 4B). However, the converse is not true: not everyone who became infected early in the outbreak is a superspreader. Individuals with many interpersonal encounters also tend to be superspreaders in the Haslemere epidemic, though again the converse is not true; that is, not all superspreaders have many encounters. Using the location data, we calculate the total number of encounters for each user, where an encounter is defined as the first time two users pass within 20 metres of each other. Fig. 4C provides a histogram of the number of unique encounters per user, distinguishing between the 90th-percentile superspreaders (red) and all others (grey). The users with the most encounters (above 41) are all superspreaders. However, the distribution of encounters made by the superspreaders is broad, with some superspreaders making as few as 10 encounters. On the other hand, some users with as many as 30 encounters are not superspreaders.

One might imagine that superspreaders could be predicted more accurately by a metric that takes into account the probability of infection. Using the transmission model (see SI) and the Haslemere mobility data, it is possible to calculate the expected number of people that a given participant in the Haslemere epidemic would directly infect if everyone else remained susceptible otherwise. We call this the user’s ‘individual reproduction number’, denoted *v*_*j*_ for user *j*, following Lloyd-Smith et al. [35]. The distribution of *v*_*j*_ for all users is depicted in Fig. 4D, again separated into the 90th-percentile superspreaders of the Haslemere epidemic (red) and all others (grey). As with the distribution of raw encounters, users with the highest *v*_*j*_ are superspreaders, but many superspreaders have low and even below-average *v*_*j*_, suggesting that this too is an unreliable way of predicting superspreaders. Superspreaders are partly determined by the stochastic course of the outbreak itself and, in real outbreak settings, possibly by individual host-pathogen dynamics [42], and so superspreaders cannot in general be predicted accurately *a priori*.

### Comparing rival notions of *R*_0_

Next, we use the Haslemere dataset to illustrate how dynamic population structure can influence empirical estimates of the basic reproduction number *R*_0_. We use an extended version of the model that was used to generate the Haslemere epidemic, for which recovery is possible (thus becoming an SEIR model rather than an SEI model), random index cases are considered, and the data are temporally looped to allow for epidemics that last beyond three days. Full details are given in the SI.

The basic reproduction number (*R*_0_) is a value of fundamental importance in infectious disease epidemiology. Defined as the expected number of secondary infections caused by a typical infectious individual introduced into a completely susceptible population [43], *R*_0_ quantifies the infectiousness of a disease in a given population, and is related to the expected final size of an outbreak and to the fraction of the population that would need to be vaccinated to prevent an outbreak from happening [44]. The mean individual reproduction number, 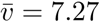, gives a model-based estimate of the basic reproduction number for the Haslemere outbreak, in that it measures the expected number of secondary infections caused by a randomly chosen individual, if the rest of the population were kept artificially susceptible [35].

Alternatively, there are many ways to empirically estimate *R*_0_ from incidence data alone. An often-used method is based on the initial growth rate of the cumulative incidence during an outbreak [45]; another is based on the outbreak’s final size [46] (see SI). Both of these empirical methods assume that the underlying population is “well-mixed”, which clearly does not hold for the population of Haslemere. We ask whether these methods can nevertheless provide accurate estimates of *R*_0_.

Fig. 5 depicts the distribution of *R*_0_ estimates from 1000 simulated outbreaks using the initial growth rate method (red) and the final size method (blue). Also depicted is the distribution of individual reproduction numbers *v*_*j*_ (black). Though the mean values of the distributions differ somewhat due to their spread (mean initial growth *R*_0_ = 4.0; mean final size *R*_0_ = 3.0; 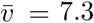), the modes of the distributions align remarkably well. However, accurate initial growth and final size *R*_0_ estimates rely on accurate knowledge of the total population size and the disease’s generation interval, respectively. For the Haslemere simulation, the size of the population is known exactly, and the generation interval may be measured directly using the span of time between each infection and its ‘parent’ (here, we set the generation interval to be the median of these time spans). In a real outbreak setting, these values may be uncertain, profoundly affecting estimates of *R*_0_. For populations with spatial or network-type structure, specifying *R*_0_ is challenging [47, 48], and the task is made even more complicated when that structure changes over time [49]. The Haslemere dataset and others like it may contribute to these discussions by allowing researchers to more precisely quantify how uncertainty in fundamental parameters propagates to estimates of *R*_0_ in the context of realistic population structures.

**Figure 5:**
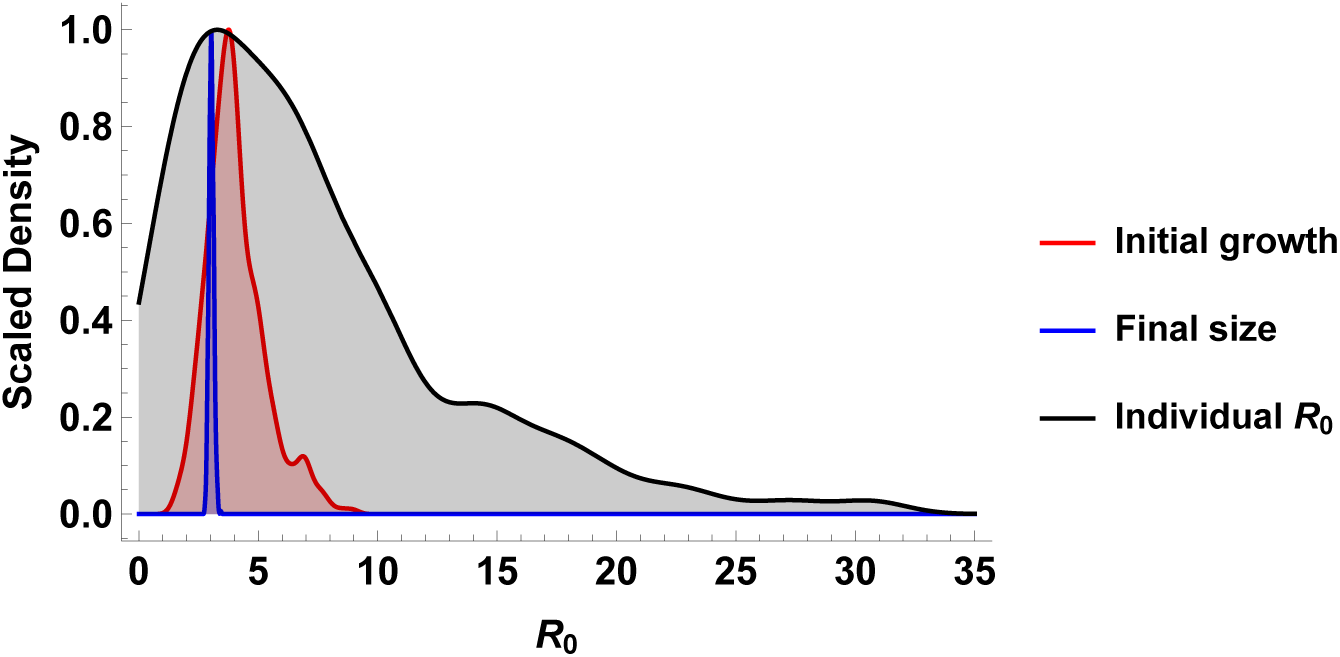
Distribution of *R*_0_ as estimated using the initial growth formula (red, mean = 4.0) and the final size formula (blue, mean = 3.0) from 1,000 simulated outbreaks. Also depicted is the smoothed distribution of individual reproduction numbers, *v*_*j*_ (black, mean = 7.3; see also Fig. 4D).

## 4 Discussion

This article presents a novel open-source dataset that captures human interaction patterns at an unprecedented scope and level of spatiotemporal detail. Previous related datasets have been restricted to relatively controlled settings such as schools [3] and community gatherings [4] or to a particular sub-population (e.g. students [40, 50, 51]), while the Haslemere dataset captures interactions for a subset of the population across an entire town for three consecutive days. The Haslemere dataset’s temporal structure in particular makes it a valuable supplement to the contact surveys already used widely in epidemiological models [52, 53]. The Haslemere dataset is limited in that it does not include the interaction patterns of children under the age of 13, who are frequently implicated as important transmitters of respiratory disease [54, 55, 56]. Using GPS to measure interpersonal distances also means that physical barriers that could obstruct disease transmission are not accounted for. We believe these limitations are a reasonable trade-off for a dataset that is freely available (precluding the inclusion of data from children) and that provides distance measurements on a continuum while minimising the barrier to participation (using mobile phones, rather than custom proximity sensors).

Though mobile phone geo-tracking capabilities have been leveraged to study human mobility before [7, 8, 9, 10, 11, 12, 13, 14], the spatial resolution in those studies is usually too coarse to characterise interpersonal interactions, and/or the data have generally not been made publicly available. Most of these studies have focused on characterising bulk movements, whereas the Haslemere dataset enhances our ability to study fine-scale interactions between individuals as they occur across time.

The need to better understand the interplay between human mobility and disease transmission has long been recognised [57]. Proxies of human movement have been integrated into disease models with some success [58, 28, 59], but the advent of wearable sensing technology, like the GPS trackers included in most smartphones, is revolutionising our insight into the human-behavioural element of disease transmission. Mobility patterns affect not only the geographic spread patterns of disease [8], but also the intensity of outbreaks [60] and the genetic diversity of circulating pathogens [25]. While the characterisation of supposedly universal laws that govern human mobility has received considerable interest [61, 62, 14], it seems likely that a close attention to the rich variety of human movement patterns, across cultures, times, and geographic scales, will yield the greatest new behavioural and epidemiological insights.

Our analysis of an epidemiological model based on the Haslemere dataset indicates that reliably predicting an outbreak’s superspreaders and estimating its basic reproduction number remain challenging tasks. In time, the availability of detailed mobility data will help address these gaps, as well as to shed light on more fundamental questions, such as: what is the most epidemiologically relevant way to delineate a population? How can the spatiotemporal dynamics of human interactions best be captured mathematically? To what extent does *a priori* knowledge of an outbreak’s superspreaders, *R*_0_, and other key parameters facilitate its control? One of the clearest potential contributions of the Haslemere dataset and others like it will be the ability to evaluate intervention strategies under more realistic conditions. Importantly, however, the Haslemere dataset does not account for behaviour changes that might occur in the context of a real outbreak. Different studies have found that social distancing during an outbreak can either dampen [63, 64] or exacerbate [65] transmission, underscoring the need for further investigation.

While the collection of human mobility data shows substantial promise for improving our understanding of disease dynamics, there remain major logistical and ethical challenges [66, 67]. The collection of the Haslemere dataset was made possible by a special collaboration between the media, academic scientists, and the broader public. The study was made possible by the clout of the BBC and by the enthusiasm of the local press, museum, and public. Even under these arguably ideal circumstances, we were only able to collect three days’ worth of data, which is too short of a duration to simulate a realistic influenza outbreak in a town. To maintain the anonymity of the volunteers, we aggregated the temporal data into five-minute bins and omitted nighttime observations. In the future, we recommend that similar studies collect data over longer time frames (on the order of weeks, if possible), that they restrict observations to every five minutes, and that they do not record nighttime movements. This should help strike a balance between relevance for infectious disease modelling and maintenance of the participants’ privacy. The dataset is also limited in that the GPS-mediated location tags from the phone’s operating system do not capture epidemiologically relevant physical barriers between individuals, such as walls. This could be surmounted for example by using wearable wireless proximity sensors as in [50], but at the significant cost of requiring volunteers to use a piece of technology not already integrated into their everyday lives. Distributing, maintaining, and recovering the requisite sensors would raise the barrier for participation, likely reducing the number of volunteers. We note that the collection of voluntary data, like the Haslemere dataset, requires more planning and oversight than the collection of so-called “convenience” datasets used in some previous mobile phone-based studies [10, 4, 58]), but it has the benefit of being more ethically straightforward, since all participants give explicit approval for the use of their data. As further studies reveal the value of the Haslemere dataset, we hope that a strong case can be made for the collection of similar data across diverse geographic settings, and that this will contribute not just to epidemiology, but to human behavioural sciences as a whole.

## 5 Materials and Methods

### Network measurements

The degree of a node is the number of links that issue from it. The local clustering coefficient for a given node is the fraction of that node’s neighbours that are also connected to one another with a link [39].

The volatility of a node *j*, denoted 1 *– γ*_*j*_ by Clauset and Eagle (2012) [40], is related to the correlation between the adjacency matrices specified by two snapshots of a dynamic network. The correlation is only calculated between elements of the adjacency matrices that are nonzero for at least one of the two snapshots, to avoid spurious correlations between the zero elements in the matrices, which are often sparse. The link persistence is defined as

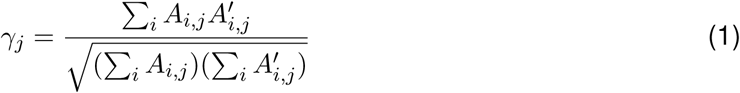

where *A* and *A’* are (unweighted) adjacency matrices corresponding to two subsequent network snapshots. We define *γ*_*j*_ = 0 if node *j* has no neighhbours in either *A* or *A’*. Link persistence is bounded between 0 and 1. The volatility for node *j* is 1 *– γ*_*j*_.

### The Haslemere epidemic model

For the Haslemere epidemic, individuals may progress from susceptible to exposed to infected (an SEI model). The exposure period lasts for 25 minutes (five time steps), after which the individual becomes infectious. There is no recovery; the epidemic is assumed to end at the end of the third day, when data collection concludes. At each time point, the force of infection *λ*_*i,j*_ between individuals *i* and *j* is modelled using an exponential curve with a cutoff:

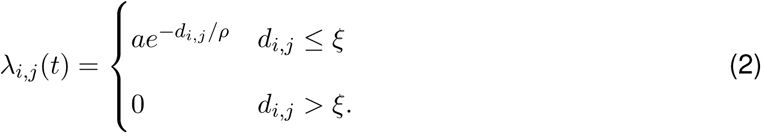

Here, *d*_*i,j*_ is the real distance in metres between individuals *i* and *j* at time *t, a* is the amplitude of the kernel, *ρ* is the decay rate, and *ξ* is the cutoff distance, after which the force of infection drops to zero. For the simulations, the parameter values are *a* = 1, *ρ* = 10 metres, and *ξ* = 20 metres (with associated kernel depicted in Fig. S1). The simulation results were shared with the study participants in a public event immediately following the end of data collection, hence a fast decision on the model and parameters was necessary (within a few hours) and sensitivity analysis was limited. For the Haslemere outbreak, we identify the most likely infector as the geographically nearest infectious individual at the time of infection. This was for convenience; in subsequent simulations, such as when defining the vaccination strategies based on the number of secondary infections, we assign infectors randomly, with probability proportional to the force of infection contributed by each possible infector at the time of infection.

The probability of infection per unit time (5 minutes) is

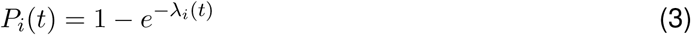

following [68, 69].

### The general epidemic model

To investigate *R*_0_ estimation methods and vaccination strategies, we define a more general epidemic model that allows for recovery and loops through the three days of available data to allow for longer epidemics. This SEIR-type model uses Eq. 2 as the transmission kernel. Individuals recover and are immune to further infection after being infected for three days (576 time steps).

## Acknowledgements

We thank all those in Haslemere who took part in the *BBC Pandemic* study. We thank Adam Kucharski for his help in specifying the project, and for his assistance in developing the ideas presented in this article. We thank Hannah Fry for the interesting discussions regarding the final model. We thank 360 Production, especially Danielle Peck and Cressida Kinnear, for making possible the collection of the dataset that underlies this work.

